## Supplementary Figures 1-5 and Supplementary Tables 1-3 for "Gut microbiome remains stable following COVID-19 vaccination in healthy and immuno-compromised individuals"

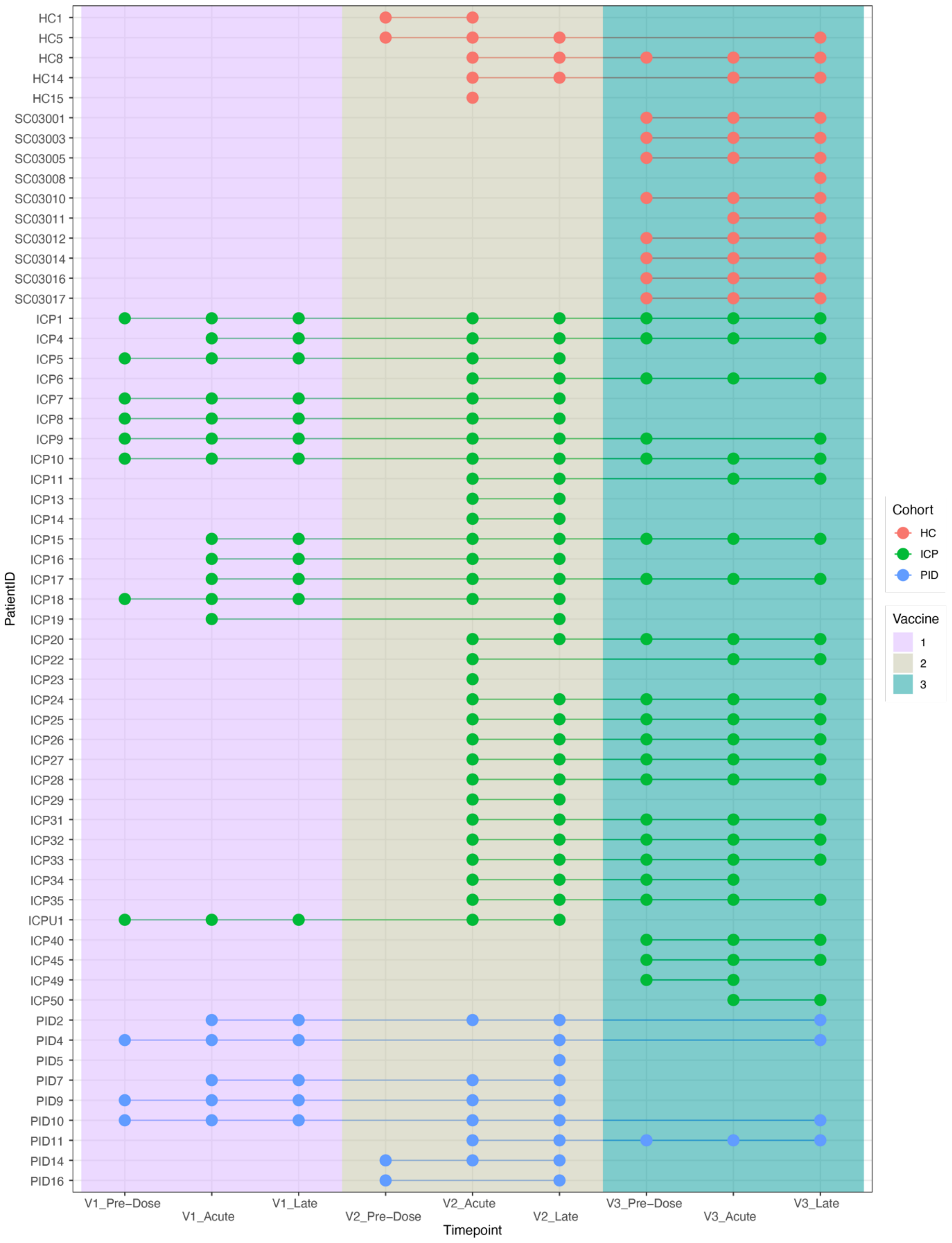


**Supplementary Figure 1.** *Participant gut microbiome sampling across three doses of COVID-19 vaccinations (V1, vaccine dose 1; V2, vaccine dose 2; V3, vaccine dose 3) from the different cohorts: healthy control (HC), immune-checkpoint therapy treated cancer patients (ICP), or patients with primary immunodeficiencies (PID).*

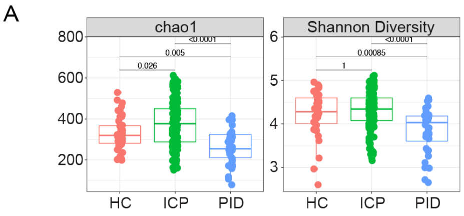


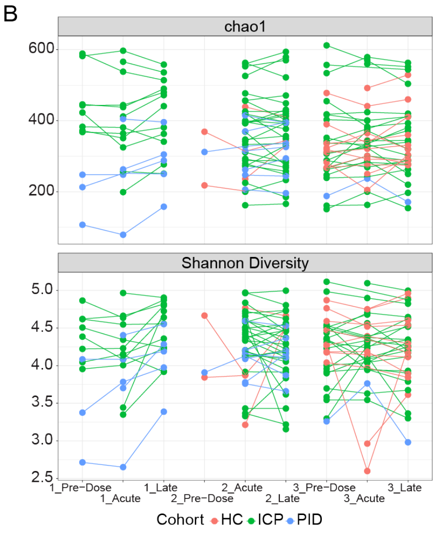


C

| | PC1 | PC2 | PC3 | PC4 | PC5 |
|---|---|---|---|---|---|
| **HC** | 0.485 | 0.960 | 0.987 | 0.473 | 0.816 |
| **ICP** | 0.776 | 0.885 | 0.481 | 0.989 | 0.890 |
| **PID** | 0.631 | 0.378 | 0.269 | 0.293 | 0.534 |

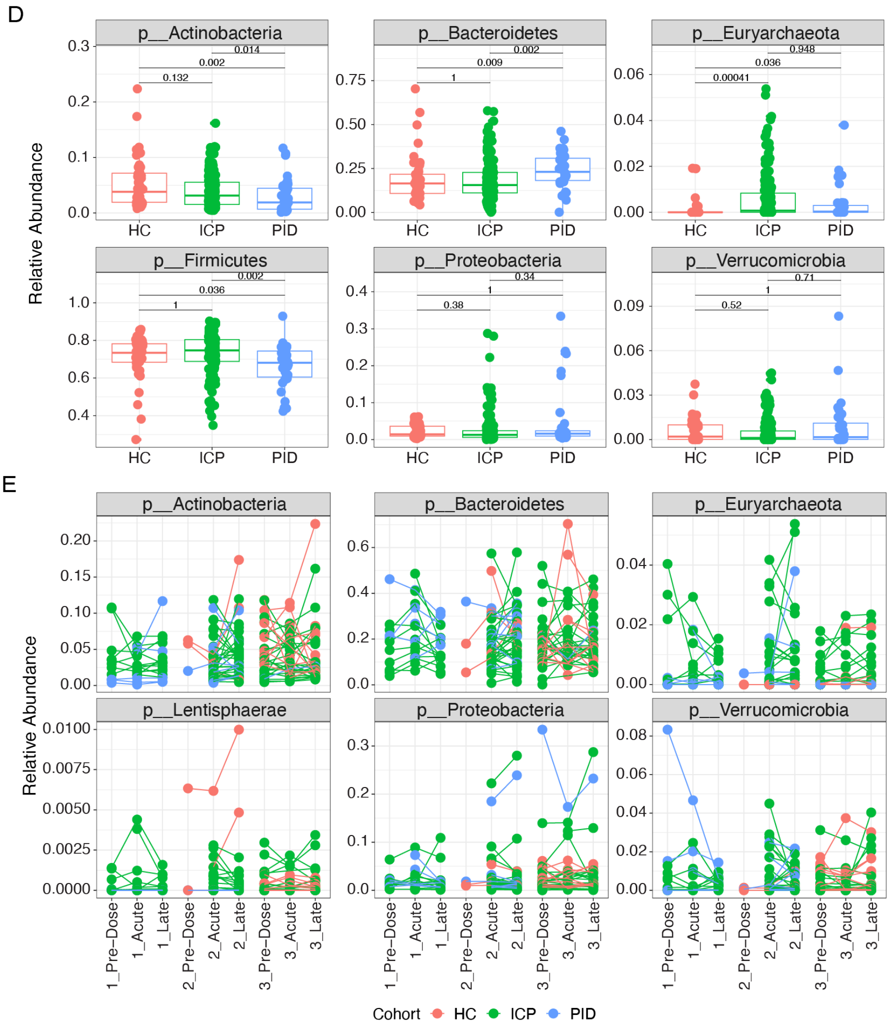


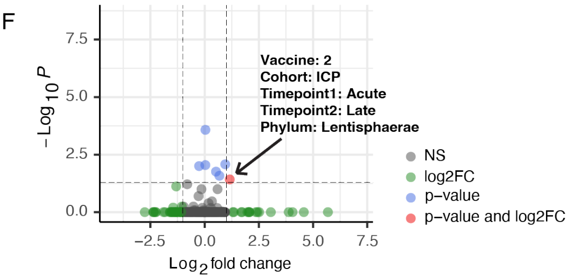

**Supplementary Figure 2.** Gut microbiome compositional differences are evident between cohorts but not vaccine timepoints.

(A) Alpha diversity measures of chao1 and Shannon diversity in samples from different cohorts*, healthy control (HC), immune-checkpoint therapy treated cancer patients (ICP), or patients with primary immunodeficiencies (PID) (N = 43 HC, 160 ICP, and 36 PID.* (B) Paired analysis of the alpha diversity measures of patient samples taken from different vaccine timepoints from each of our cohorts *(N = 40 HC, 153 ICP, and 31 PID)*. (C) Reported p values of the linear model comparing baseline model of fixed patient effects on the explained variance, to the model utilising the vaccine timepoints of the patient samples as random effects when using the principal components (PC) (healthy controls (HCs), immune checkpoint treated cancer patients (ICP), Primary Immunodeficient patients (PID)). (D) Relative abundance of the 6 most prevalent phyla in our patient samples from within each of our patient cohorts and separated by the vaccine timepoint from which the sample was taken *(N = 43 HC, 160 ICP, and 36 PID).* (E) Paired analysis of the 6 most prevalent phyla in our patient samples taken from different vaccine timepoints from each of our cohorts *(N = 40 HC, 153 ICP, and 31 PID)*. (F) Volcano plot of the paired relative phylum abundance between two timepoints across all pair combinations unique to different vaccine doses and different cohorts. Colours represent the significance indicated in the legend. Statistical testing within figures was performed using Wilcoxon test and adjusted for multiple testing using bonferonni, paired where appropriate.

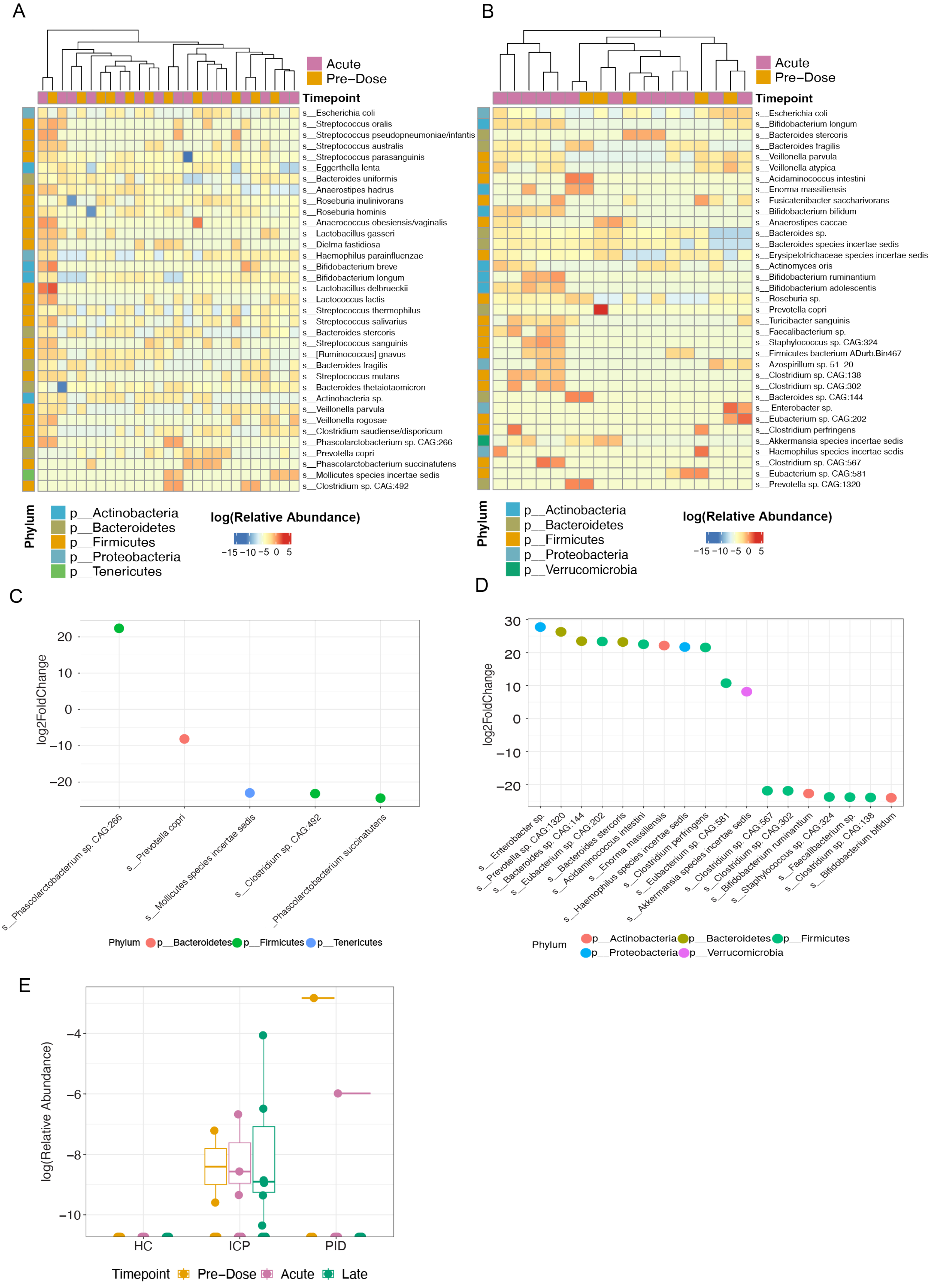
A
Acute
Pre−Dose
Timepoint
s__Escherichia coli
s__Streptococcus oralis
s__Streptococcus pseudopneumoniae/infantis
s__Streptococcus australis
s__Streptococcus parasanguinis
s__Eggerthella lenta
s__Bacteroides uniformis
s__Anaerostipes hadrus
s__Roseburia inulinivorans
s__Roseburia hominis
s__Anaerococcus obesiensis/vaginalis
s__Lactobacillus gasseri
s__Dielma fastidiosa
s__Haemophilus parainfluenzae
s__Bifidobacterium breve
s__Bifidobacterium longum
s__Lactobacillus delbrueckii
s__Lactococcus lactis
s__Streptococcus thermophilus
s__Streptococcus salivarius
s__Bacteroides stercoris
s__Streptococcus sanguinis
s__[Ruminococcus] gnavus
s__Bacteroides fragilis
s__Streptococcus mutans
s__Bacteroides thetaiotaomicron
s__Actinobacteria sp.
s__Veillonella parvula
s__Veillonella rogosae
s__Clostridium saudiense/disporicum
s__Phascolarctobacterium sp. CAG:266
s__Prevotella copri
s__Phascolarctobacterium succinatutens
s__Mollicutes species incertae sedis
s__Clostridium sp. CAG:492
Phylum
p__Actinobacteria
p__Bacteroidetes
p__Firmicutes
p__Proteobacteria
p__Tenericutes
log(Relative Abundance)
−15 −10 −5 0 5
B
Acute
Pre−Dose
Timepoint
s__Escherichia coli
s__Bifidobacterium longum
s__Bacteroides stercoris
s__Bacteroides fragilis
s__Veillonella parvula
s__Veillonella atypica
s__Acidaminococcus intestini
s__Enorma massiliensis
s__Fusicatenibacter saccharivorans
s__Bifidobacterium bifidum
s__Anaerostipes caccae
s__Bacteroides sp.
s__Bacteroides species incertae sedis
s__Erysipelotrichaceae species incertae sedis
s__Actinomyces oris
s__Bifidobacterium ruminantium
s__Bifidobacterium adolescentis
s__Roseburia sp.
s__Prevotella copri
s__Turicibacter sanguinis
s__Faecalibacterium sp.
s__Staphylococcus sp. CAG:324
s__Firmicutes bacterium ADurb.Bin467
s__Azospirillum sp. 51_20
s__Clostridium sp. CAG:138
s__Clostridium sp. CAG:302
s__Bacteroides sp. CAG:144
s__Enterobacter sp.
s__Eubacterium sp. CAG:202
s__Clostridium perfringens
s__Akkermansia species incertae sedis
s__Haemophilus species incertae sedis
s__Clostridium sp. CAG:567
s__Eubacterium sp. CAG:581
s__Prevotella sp. CAG:1320
Phylum
p__Actinobacteria
p__Bacteroidetes
p__Firmicutes
p__Proteobacteria
p__Verrucomicrobia
log(Relative Abundance)
−15 −10 −5 0 5
C
log2FoldChange
20
10
0
−10
−20
s__Phascolarctobacterium sp. CAG:266
s__Prevotella copri
s__Mollicutes species incertae sedis
s__Clostridium sp. CAG:492
s__Phascolarctobacterium succinatutens
Phylum p__Bacteroidetes p__Firmicutes p__Tenericutes
D
log2FoldChange
30
20
10
0
−10
−20
s__Enterobacter sp.
s__Prevotella sp. CAG:1320
s__Bacteroides sp. CAG:144
s__Eubacterium sp. CAG:202
s__Bacteroides stercoris
s__Acidaminococcus intestini
s__Enorma massiliensis
s__Haemophilus species incertae sedis
s__Clostridium perfringens
s__Eubacterium sp. CAG:581
s__Akkermansia species incertae sedis
s__Clostridium sp. CAG:567
s__Clostridium sp. CAG:302
s__Bifidobacterium ruminantium
s__Staphylococcus sp. CAG:324
s__Faecalibacterium sp.
s__Clostridium sp. CAG:138
s__Bifidobacterium bifidum
Phylum p__Actinobacteria p__Bacteroidetes p__Firmicutes p__Proteobacteria p__Verrucomicrobia
E
log(Relative Abundance)
−4
−6
−8
−10
HC
ICP
PID
Timepoint Pre−Dose Acute Late

**Supplementary Figure 3.** Differential abundance analysis of HC and PID cohort samples taken at pre-dose and acutely after COVID-19 vaccination.

(A) DESeq analysis of HC cohort (N = 11 Pre-Dose, 16 Acute) and PID cohort (N = 5 Pre-Dose, 13 Acute) (B), along with corresponding log2FoldChange of significantly different bacterial species between pre-dose and acute samples in HC samples (C) and PID samples (D), colours represent different phyla. (E) Relative abundance of Enterobacter sp. in our cohorts.

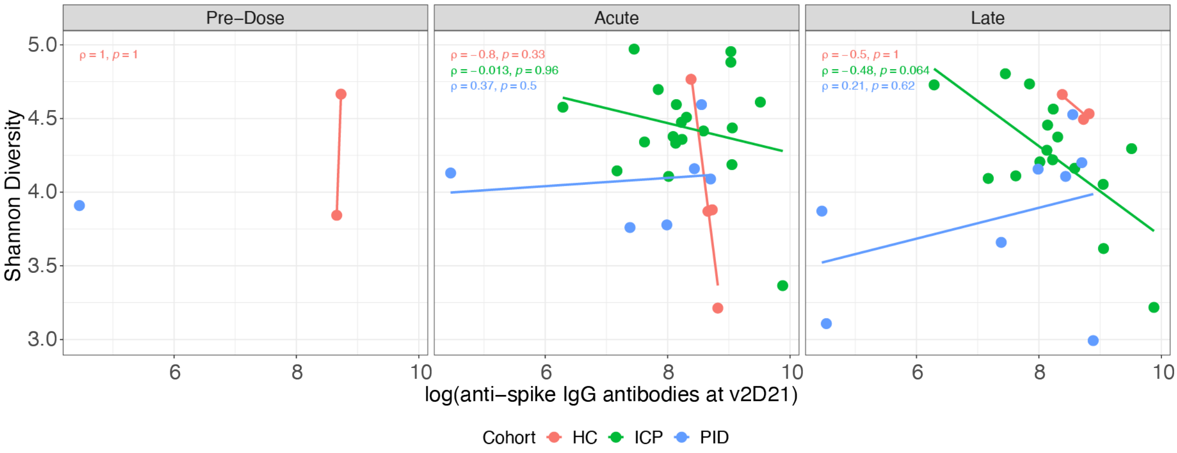


**Supplementary Figure 4.** Absence of correlation between vaccine efficacy and gut microbiome composition. Second dose *anti-spike IgG antibody levels assessed against Shannon diversity of fecal samples; each point represents a different sample taken at one of the three vaccine timepoints. Colours represent cohorts, within healthy control (HC), immune-checkpoint therapy treated cancer patients (ICP) and patients with primary immunodeficiencies (PID. rho and p values from Spearman correlation testing displayed (N = 9 HC, 35 ICP, and 15 PID).*

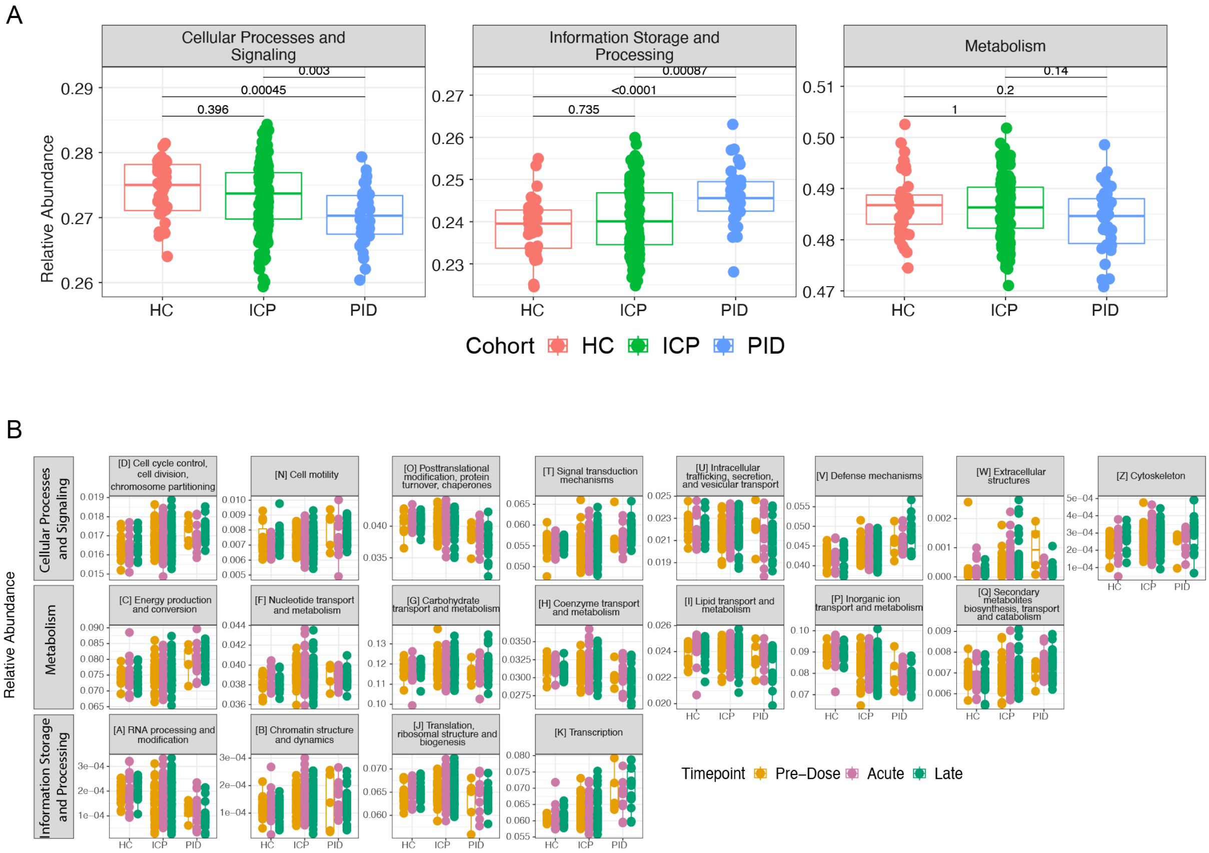


Supplementary Figure 5. Functional annotation differences between cohorts, but not between vaccine timepoints.

(A) *The relative abundance of the highest functional annotation level within patient samples from different patient cohort (healthy controls, HC, immune checkpoint therapy treated cancer patients, ICP, and primary immunodeficient patients PID), separated by the vaccine timepoints from which the sample was taken. (B) Relative abundance of the remaining 19 out of a possible 22 functional annotations in our patient samples from within each of our patient cohorts and separated by the vaccine timepoint from which the sample was taken. Statistical testing performed using Wilcoxon test and adjusted for multiple testing using FDR (N = 43 HC, 160 ICP, and 36 PID).*

| Diversity_Measure | Vaccine | Cohort | Timepoint_1 | Timepoint_2 | Samples | P_value | padj |
|---|---|---|---|---|---|---|---|
| chao1 | 1 | ICP | Pre−Dose | Acute | 8 | 0.107327557761612 | 0.6439653 |
| chao1 | 1 | ICP | Pre−Dose | Late | 8 | 0.833634883024682 | 1.0000000 |
| chao1 | 1 | ICP | Acute | Late | 12 | 0.84451926747294 | 1.0000000 |
| chao1 | 1 | PID | Pre−Dose | Acute | 3 | 1 | 1.0000000 |
| chao1 | 1 | PID | Pre−Dose | Late | 3 | 0.18144920772142 | 1.0000000 |
| chao1 | 1 | PID | Acute | Late | 5 | 0.787406490666269 | 1.0000000 |
| diversity_shannon | 1 | ICP | Pre−Dose | Acute | 8 | 0.362726506485098 | 1.0000000 |
| diversity_shannon | 1 | ICP | Pre−Dose | Late | 8 | 0.141482121482793 | 0.8488927 |
| diversity_shannon | 1 | ICP | Acute | Late | 12 | 0.224015384088612 | 1.0000000 |
| diversity_shannon | 1 | PID | Pre−Dose | Acute | 3 | 0.789268026134281 | 1.0000000 |
| diversity_shannon | 1 | PID | Pre−Dose | Late | 3 | 0.18144920772142 | 1.0000000 |
| diversity_shannon | 1 | PID | Acute | Late | 5 | 0.589638551626567 | 1.0000000 |
| chao1 | 2 | HC | Acute | Late | 3 | 0.789268026134281 | 1.0000000 |
| chao1 | 2 | ICP | Acute | Late | 28 | 0.674165890945232 | 1.0000000 |
| chao1 | 2 | PID | Acute | Late | 7 | 0.932646638965876 | 1.0000000 |
| diversity_shannon | 2 | HC | Acute | Late | 3 | 0.422678074170635 | 0.8453561 |
| diversity_shannon | 2 | ICP | Acute | Late | 28 | 0.0634719464247325 | 0.1269439 |
| diversity_shannon | 2 | PID | Acute | Late | 7 | 0.554113130069446 | 1.0000000 |
| chao1 | 3 | HC | Pre−Dose | Acute | 9 | 0.635586122278767 | 1.0000000 |
| chao1 | 3 | HC | Pre−Dose | Late | 9 | 0.342828839723404 | 1.0000000 |
| chao1 | 3 | HC | Acute | Late | 11 | 0.449803788345075 | 1.0000000 |
| chao1 | 3 | ICP | Pre−Dose | Acute | 20 | 0.155895425145994 | 0.9353726 |
| chao1 | 3 | ICP | Pre−Dose | Late | 19 | 0.601249670135047 | 1.0000000 |
| chao1 | 3 | ICP | Acute | Late | 21 | 0.87570099927198 | 1.0000000 |
| diversity_shannon | 3 | HC | Pre−Dose | Acute | 9 | 0.0972010883062347 | 0.5832065 |
| diversity_shannon | 3 | HC | Pre−Dose | Late | 9 | 0.0580240199462214 | 0.3481441 |
| diversity_shannon | 3 | HC | Acute | Late | 11 | 0.504879873834092 | 1.0000000 |
| diversity_shannon | 3 | ICP | Pre−Dose | Acute | 20 | 0.640744318210405 | 1.0000000 |
| diversity_shannon | 3 | ICP | Pre−Dose | Late | 19 | 0.732306833608737 | 1.0000000 |
| diversity_shannon | 3 | ICP | Acute | Late | 21 | 0.754418293600069 | 1.0000000 |

*Supplementary Table 1. Paired sample alpha diversity analysis from different patient cohorts (healthy controls, HC, immune checkpoint therapy treated cancer patients, ICP, and primary immunodeficient patients PID). Statistical testing using paired Wilcoxon test, with bonferonni adjustment for multiple testing.*

```
permanova <- adonis2(t(otu) ~ Patient*Vaccine*Timepoint + Sex + Age,
data = meta, permutations=999, method = "bray")
```

| | Df | SumOfSqs | R2 | F | Pr(>F) |
|---|---|---|---|---|---|
| *Patient* | 58 | 5.25969000530358 | 0.688469968284202 | 4.5975033754811 | 0.003 |
| *Vaccine* | 2 | 0.0348401851718894 | 0.00456042488361858 | 0.883161972338744 | 0.662 |
| *Timepoint* | 2 | 0.0202817693779327 | 0.00265479317341762 | 0.514121476621395 | 0.89 |
| *Patient:Vaccine* | 43 | 0.859041502723111 | 0.112444702166522 | 1.01282775158248 | 0.605 |
| *Patient:Timepoint* | 92 | 1.03439397364817 | 0.135397558698865 | 0.570017325530319 | 0.892 |
| *Vaccine:Timepoint* | 3 | 0.0598522755605799 | 0.00783439598443623 | 1.01146140053035 | 0.573 |
| *Patient:Vaccine:Timepoint* | 36 | 0.3321305647264 | 0.0434744099239363 | 0.467731097120218 | 0.952 |
| *Residual* | 2 | 0.0394493719873687 | 0.00516374688500445 | NA | NA |
| *Total* | 238 | 7.63967964849902 | 1 | NA | NA |

*Supplementary Table 2. PERMANOVA analysis describing the influence of study variables on the variance seen in the composition of the gut microbiome samples from our patients.*

| Vaccine | Cohort | Timepoint_1 | Timepoint_2 | Phylum | foldchange | comparisons | P_value | padj | log2FC |
|---|---|---|---|---|---|---|---|---|---|
| 2 | ICP | Acute | Late | p__Firmicutes | 1.02569986537809 | 28 | 1.76741562992507e−05 | 0.0002651123 | 0.03660864 |
| 2 | ICP | Acute | Late | p__Verrucomicrobia | 1.9317442888861 | 28 | 0.000556247323105197 | 0.0083437098 | 0.94990413 |
| 3 | ICP | Acute | Late | p__Firmicutes | 1.02448433089915 | 21 | 0.00019995878734921 | 0.0089981454 | 0.03489792 |
| 2 | ICP | Acute | Late | p__Bacteroidetes | 0.837832989721559 | 28 | 0.000663271058611996 | 0.0099490659 | −0.25526540 |
| 3 | ICP | Acute | Late | p__Bacteria phylum incertae sedis | 1.43802342229653 | 21 | 0.00038470759403593 | 0.0173118417 | 0.52408717 |
| 3 | ICP | Pre−Dose | Late | p__Bacteria phylum incertae sedis | 1.60364273027069 | 19 | 0.000584658962712894 | 0.0263096533 | 0.68135276 |
| 2 | ICP | Acute | Late | p__Lentisphaerae | 2.23005380142644 | 28 | 0.00252617426850217 | 0.0378926140 | 1.15707852 |

*Supplementary Table 3. Paired sample analysis of the relative abundance of phylum from different patient cohorts (healthy controls, HC, immune checkpoint therapy treated cancer patients, ICP, and primary immunodeficient patients PID). Statistical testing using paired Wilcoxon test, with bonferonni adjustment for multiple testing; log 2 fold-change (log2FC). Only those values with padj < 0.05 are shown.*
